# Chromosome-scale genome assembly of bread wheat’s wild relative *Triticum timopheevii*

**DOI:** 10.1101/2024.01.16.575864

**Authors:** Surbhi Grewal, Cai-yun Yang, Duncan Scholefield, Stephen Ashling, Sreya Ghosh, David Swarbreck, Joanna Collins, Eric Yao, Taner Z. Sen, Michael Wilson, Levi Yant, Ian P. King, Julie King

## Abstract

Wheat (*Triticum aestivum*) is one of the most important food crops with an urgent need for increase in its production to feed the growing world. *Triticum timopheevii* (2n = 4x = 28) is an allotetraploid wheat wild relative species containing the A^t^ and G genomes that has been exploited in many pre-breeding programmes for wheat improvement. In this study, we report the generation of a chromosome-scale reference genome assembly of *T. timopheevii* accession PI 94760 based on PacBio HiFi reads and chromosome conformation capture (Hi-C). The assembly comprised a total size of 9.35 Gb, featuring a contig N50 of 42.4 Mb, and 166,325 predicted gene models. DNA methylation analysis showed that the G genome had on average more methylated bases than the A^t^ genome. The G genome was also more closely related to the S genome of *Aegilops speltoides* than to the B genome of hexaploid or tetraploid wheat. In summary, the *T. timopheevii* genome assembly provides a valuable resource for genome-informed discovery of agronomically important genes for food security.

## Background and Summary

The *Triticum* genus comprises many wild and cultivated wheat species including diploid, tetraploid and hexaploid forms. The polyploid species originated after hybridisation between *Triticum* and the neighbouring Aegilops genus (goatgrass). The tetraploid species, *Triticum turgidum* (2n = 4x = 28, AABB), also known as emmer wheat, and *Triticum timopheevii* (2n = 4x = 28, A^t^A^t^GG) are polyphyletic. *Triticum urartu* Thum. ex Gandil (2n = 2x = 14, AA) is the A genome donor for both these species^1^ whereas, the B and G genomes are closely related to the S genome of *Aegilops speltoides*^2^. Both tetraploid species have wild and domesticated forms, i.e., *T. turgidum* L. ssp. *dicoccoides* (Körn. ex Asch. & Graebn.) Thell. and ssp. *dicoccum* (Schrank ex Schübl.) Thell., respectively, and *T. timopheevii* (Zhuk.) Zhuk. ssp. *armeniacum* (Jakubz.) Slageren and ssp. timopheevii, respectively. Additionally, tetraploid durum wheat *T. turgidum* L. ssp. *durum* (Desf.) Husn. (2n = 4x = 28, AABB), used for pasta production, and hexaploid bread wheat *Triticum aestivum* L. (2n = 6x = 42, AABBDD) evolved from domesticated emmer wheat with the latter originating through hybridisation with *Aegilops tauschii* (D genome donor) 6,000-7,000 years ago. Hexaploid *Triticum zhukovskyi* (AAGGA^m^A^m^) originated from hybridisation of cultivated T. timopheevii and cultivated einkorn *Triticum monococcum*^3^ (2n = 2x = 14, A^m^A^m^).

The G genome is only found in *T. timopheevii* and *T. zhukovskyi* and is virtually identical to the S genome on a molecular level^4,5^ but differs from it, and the B genome, due to a number of chromosomal rearrangements and translocations involving the A^t^ genome^6^. The most studied are the 6A^t^/1G/4G and 4G/4A^t^/3A^t^ translocations in *T. timopheevii*^7-10^.

*Triticum timopheevii* ssp. *timopheevii* has been exploited in various studies for wheat improvement as it has been shown to be an abundant source for genetic variation for many traits such as resistance to leaf rust^11-13^, stem rust^14-16^, powdery mildew^16-18^, fusarium head blight^19,20^ Hessian fly, Septoria blotch, wheat curl mite and tan spot^21^. It has also been shown to have tolerance to abiotic stresses such as salinity^22,23^ and be a good source for traits affecting grain quality such as milling yield and grain protein^24^ and grain mineral content^25^. During sequence analysis of reference quality assemblies (RQA) of 10 wheat cultivars, recent studies found two of them, cv. LongReach Lancer and cv. Julius, contained major introgressions on Chr2B (among others) potentially originating from *T. timopheevii*^26,27^. Introgressions from *T. timopheevii* have also been found in many other wheat accessions present in genebank collections^28^. Pre-breeding programmes involving the introgression of the whole genome of *T. timopheevii*, in small segments, into bread wheat^10,29^ with diagnostic KASP markers that can track these introgressions in wheat^29,30^ have provided promising new germplasm and tools to the wheat research community.

In this study, we report a chromosome-scale reference genome sequence assembly for *T. timopheevii* by integrating chromatin conformation capture (Hi-C) derived short-reads^31^ with PacBio HiFi long-reads^32^. The assembly was annotated for gene models and repeats. CpG methylation along the chromosomes was inferred from the PacBio CCS data. Known chromosomal translocations within and between the A^t^ and G genomes were confirmed, and new chromosome rearrangements were found in comparison to wild emmer wheat. The high-quality *T. timopheevii* genome assembly obtained in this study provides a reference for the G genome of the *Triticum* genus. This new resource will form the basis to study chromosome rearrangements across different Triticeae species and will be explored to detect A^t^ and G genome introgressions in durum and bread wheat allowing future genome-informed gene discoveries for various agronomic traits.

## Methods

### Plant material, nucleic acid extraction and sequencing

High molecular weight (HMW) DNA was extracted from a young seedling (dark-treated for 48 hours) of *T. timopheevii* accession PI 94760 (United States National Plant Germplasm System, NPGS available at https://npgsweb.ars-grin.gov/gringlobal/search) using a modified Qiagen Genomic DNA extraction protocol (https://doi.org/10.17504/protocols.io.bafmibk6)^33^. DNA was sheared to the appropriate size range (15–20⍰kb) and PacBio HiFi sequencing libraries were constructed by Novogene (UK) Company Limited. Sequencing was performed on 10 SMRT cells of the PacBio Sequel II system in CCS mode with kinetics option to generate ∼267 Gb (∼28-fold coverage) of long HiFi reads (Table S1). Four Hi-C libraries were prepared using leaf samples (from the same plant used for HMW DNA extraction), at Phase Genomics (Seattle, USA) using the Proximo® Hi-C Kit for plant tissues according to the manufacturer’s protocol. The Hi-C libraries were sequenced on an Illumina NovaSeq 6000 S4 platform to generate ∼2.8 billion of paired end 150bp reads (∼842 Gb raw data; ∼89-fold coverage; Table S2).

Total RNA was extracted from seedlings (3-leaf stage), seedlings at dusk, roots, flag leaves, spikes and grains. Flag leaf and whole spike were collected at 7 days post-anthesis and whole grains were collected at 15 days post-anthesis. In brief, 100⍰mg of ground powder from each tissue was used for RNA isolation using the RNeasy Plant Mini Kit (#74904, QIAGEN Ltd UK) following manufacturer’s instructions. The RNA samples were split into 2 aliquots, one for mRNA sequencing (RNA-Seq) and one for Iso-Seq^34^. Library construction for both types of sequencing was carried out by Novogene (UK) Company Limited. Illumina NovaSeq 6000 S4 platform was used for mRNA sequencing to generate on average 450 million reads (∼67 Gb of 2 x 150bp reads) for each sample (Table S3). The second set of RNA aliquots from each of the six tissues were pooled into one sample and sequenced on the PacBio Sequel II system using the Iso-Seq pipeline to generate 4.47 Gb of Iso-Seq data (Table S4) which was analysed using PacBio Iso-Seq analysis pipeline (SMRT Link v12.0.0.177059).

Plants were grown in a glasshouse in 2L pots containing John Innes No. 2 soil and maintained at 18 – 25 °C under 16 h light and 8 h dark conditions. All sequencing was carried out by Novogene (UK) Company Limited.

### Cleaning of sequencing data

The HiFi sequencing read files in BAM format were converted and combined into one fastq file using bam2fastq v1.3.1 (available at https://github.com/jts/bam2fastq). Reads with PacBio adapters were removed using cutadapt v4.1^35^ with parameters: --error-rate=0.1 --times=3 --overlap=35 -- action=trim --revcomp --discard-trimmed. Hi-C reads were trimmed to remove Illumina adapters using Trimmomatic v0.39^36^ with parameters ILLUMINACLIP:TruSeq3-PE-2.fa:2:30:10:2:keepBothReads SLIDINGWINDOW:4:20 MINLEN:40 CROP:150.

### Genome size estimation

The size of the *T. timopheevii* genome was estimated by using k-mer (k⍰= ⍰32) distribution analysis with Jellyfish v2.2.10^37^ on the cleaned HiFi reads^38^. A k-mer count histogram was generated and the size of the *T. timopheevii* genome was estimated as ∼9.46 Gb with heterozygosity of 0.001% (Fig. 1), using GenomeScope v2.0^39^ (available at http://qb.cshl.edu/genomescope/genomescope2.0/) with parameters: ploidy = 2, k-mer length = 32, max k-mer coverage = 1000000 and average k-mer coverage = 10.

**Figure 1.**
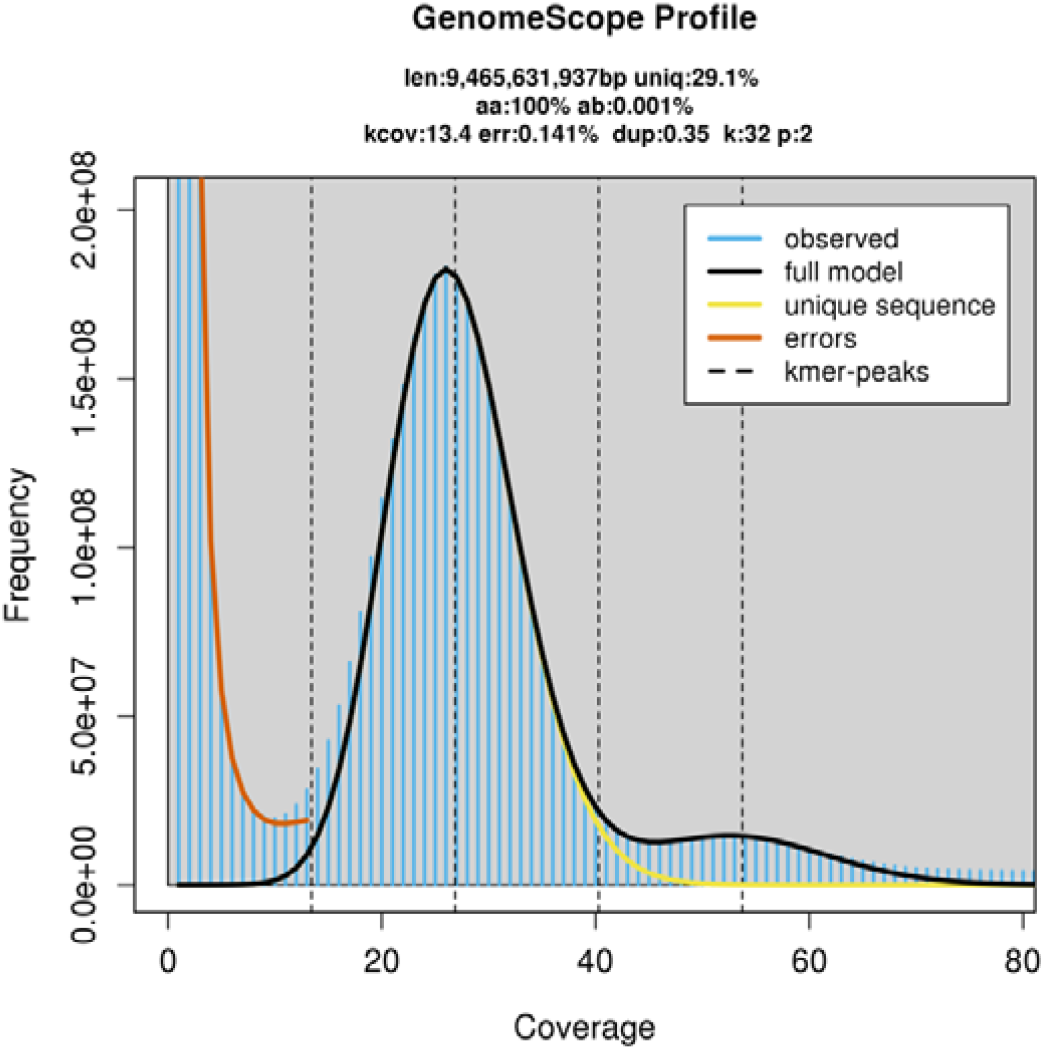
Genomescope profile for 32-mers based on HiFi reads.

### Chromosome-scale genome assembly

The cleaned HiFi reads were assembled into the initial set of contigs using hifiasm v.0.16.1^40^ with default parameters for an inbred species (-l 0) and the dataset was assessed using gfastats v1.3.1^41^. The contig assembly had a total size of ∼9.41 Gb, with a contig N50 value of 43.12⍰Mb. Genome completeness was assessed using the Benchmarking Universal Single-Copy Orthologs (BUSCO v5.3.2)^42^ program with the embryophyta_odb10 database which yielded 99% of the complete BUSCO genes. Contaminants (contigs other than those categorised as Streptophyta or no hit) were identified using BlobTools v1.1.1^43^ and removed.

To achieve chromosome-level assembly, the trimmed Hi-C data^44^ was mapped onto the decontaminated contig assembly using the Arima Genomics® mapping pipeline (available at https://github.com/ArimaGenomics/mapping_pipeline) and chromosome construction was conducted using the Salsa2^45^ pipeline (available at https://github.com/marbl/SALSA) with default parameters and GATC as the cutting site for the restriction enzyme (DpnII). The Hi-C contact map for the scaffold assembly was constructed using PretextMap v0.1.9 and the chromatin contact matrix was manually corrected using PretextView v0.2.5 by following the Rapid Curation pipeline^46^ (https://gitlab.com/wtsi-grit/rapid-curation). The curated assembly was assessed using gfastats to consist of 14 pseudomolecules and 1656 unplaced scaffolds with a total length of 9,350,839,849 bp (including gaps) and a contig N50 of 42.4 Mb (Table 1). The orientation and the chromosome name of each pseudomolecule were determined based on homology with the wheat cv. Chinese Spring assembly RefSeq2.1^47^ A and B subgenomes, using dotplot comparison of sequence alignments produced by MUMmer’s (v3.23^48^) nucmer aligner and viewed on Dot (available at https://github.com/marianattestad/dot). The pseudomolecules were thus, renamed into the 14 *T. timopheevii* chromosomes, seven A^t^ genome chromosomes with a total length of ∼4.85 Gb and consisting of 119 contigs and seven G genome chromosomes with a total length of ∼4.40 Gb and consisting of 529 contigs (Table 2).

**Table 1.** Summary statistics for genome assembly of *Triticum timopheevii*.

| Assembly characteristics | Value |
| --- | --- |
| Number of scaffolds | 1,670 |
| Total scaffold length (bp) | 9,350,839,849 |
| Scaffold N50 (bp) | 671,191,297 |
| Largest scaffold (bp) | 771,176,557 |
| No. of contigs | 2,304 |
| Total contig length (bp) | 9,350,587,949 |
| Average contig length (bp) | 4,058,415 |
| Contig N50 (bp) | 42,410,373 |
| Largest contig (bp) | 311,469,246 |
| GC content (%) | 46 |
| BUSCO evaluation (% of complete BUSCO genes) | 98.9 |

**Table 2.** Statistics of the *Triticum timopheevii* pseudomolecules.

| Chromosome | Length (bp) | Number of contigs | Number of gene models |
| --- | --- | --- | --- |
| Chr1A<sup>t</sup> | 614,431,332 | 14 | 9,982 |
| Chr1G | 495,016,746 | 50 | 8,777 |
| Chr2A<sup>t</sup> | 767,071,137 | 10 | 12,729 |
| Chr2G | 671,256,291 | 72 | 13,941 |
| Chr3A<sup>t</sup> | 670,741,101 | 10 | 9,489 |
| Chr3G | 671,191,297 | 75 | 13,452 |
| Chr4A<sup>t</sup> | 771,176,557 | 23 | 12,878 |
| Chr4G | 643,128,204 | 68 | 9,936 |
| Chr5A<sup>t</sup> | 694,350,238 | 12 | 11,821 |
| Chr5G | 641,290,954 | 78 | 13,079 |
| Chr6A<sup>t</sup> | 585,824,631 | 33 | 9,011 |
| Chr6G | 589,079,669 | 87 | 11,406 |
| Chr7A<sup>t</sup> | 745,638,687 | 17 | 12,863 |
| Chr7G | 692,654,486 | 99 | 14,851 |
| Unplaced scaffolds | 97,988,519 | 1656 | 2,110 |
| **Total** | **9,350,839,849** | **2,304** | **166,325** |

### Organellar genome assembly

*De novo* assembly of the organelle genomes was carried out using the Oatk pipeline (available at https://github.com/c-zhou/oatk) with the cleaned HiFi reads. The circular chloroplast and mitochondrial contigs were assembled with a total size of 136,158 bp and 443,464 bp, respectively. Any unanchored contigs that aligned to these extranuclear genomes were removed from the final assembly.

### Genome annotation

Gene models were generated from the *T. timopheevii* assembly using REAT - Robust and Extendable eukaryotic Annotation Toolkit (https://github.com/EI-CoreBioinformatics/reat) and Minos (https://github.com/EI-CoreBioinformatics/minos) which make use of Mikado^49^ (https://github.com/EI-CoreBioinformatics/mikado), Portcullis (https://github.com/EI-CoreBioinformatics/portcullis) and many third-party tools (listed in the above repositories). A consistent gene naming standard^50^ was used to make the gene models uniquely identifiable.

#### 1. Repeat identification

Repeat annotation was performed using EI-Repeat version 1.3.4 pipeline (https://github.com/EI-CoreBioinformatics/eirepeat) which uses third party tools for repeat calling. In the pipeline, RepeatModeler (v1.0.11 - http://www.repeatmasker.org/RepeatModeler/) was used for de novo identification of repetitive elements from the assembled *T. timopheevii* genome. High copy protein coding genes potentially included in the RepeatModeler library were identified and effectively removed by running RepeatMasker v4.0.7 using a curated set of high confidence *T. aestivum* coding genes to hard mask the RepeatModeler library; transposable element genes were first excluded from the *T. aestivum* coding gene set by running TransposonPSI (r08222010). Unclassified repeats were searched in a custom BLAST database of organellar genomes (mitochondrial and chloroplast sequences from *Triticum* in the NCBI nucleotide division). Any repeat families matching organellar DNA were also hard-masked. Repeat identification was completed by running RepeatMasker v4.0.72 with a RepBase embryophyte library and with the customized RepeatModeler library (i.e. after masking out protein coding genes), both using the - nolow setting.

#### 2. Reference guided transcriptome reconstruction

Gene models were derived from the RNA-Seq reads, Iso-Seq transcripts (122,253 high quality and 82 low quality isoforms; Supplementary File 1) and Full-Length Non-Concatamer Reads (FLNC) using the REAT transcriptome workflow. HISAT2 v2.2.1^51^ was selected as the short read aligner with Iso-Seq transcripts aligned with minimap2 v2.18-r1015^52^, maximum intron length was set as 50,000 bp and minimum intron length to 20bp. Iso-Seq alignments were required to meet 95% coverage and 90% identity. High-confidence splice junctions were identified by Portcullis v 1.2.4^53^. RNA-Seq Illumina reads were assembled for each tissue with StringTie2 v2.1.5^54^ and Scallop v0.10.5^55^, while FLNC reads were assembled using StringTie2 (Table S5). Gene models were derived from the RNA-Seq assemblies and Iso-Seq and FLNC alignments with Mikado. Mikado was run with all Scallop, StringTie2, Iso-Seq and FLNC alignments and a second run with only Iso-Seq and FLNC alignments (Table S6).

#### 3. Cross-species protein alignment

Protein sequences from 10 Poaceae species (Table S7) were aligned to the *T. timopheevii* assembly using the REAT Homology workflow with options --annotation_filters aa_len --alignment_species Angiosp --filter_max_intron 20000 -- filter_min_exon 10 --alignment_filters aa_len internal_stop intron_len exon_len splicing -- alignment_min_coverage 90 --junction_f1_filter 40 --post_alignment_clip clip_term_intron-exon - -term5i_len 5000 --term3i_len 5000 --term5c_len 36 --term3c_len 36. The REAT Homology workflow aligns proteins with spaln v2.4.7^56^ and filters and generates metrics to remove misaligned proteins. Simultaneously, the same protein set were also aligned using miniprot v0.3^57^ and similarly filtered as in the REAT homology workflow. The aligned proteins from both methods were clustered into loci and a consolidated set of gene models were derived via Mikado.

#### 4. Evidence guided gene prediction

The evidence guided annotation of protein coding genes based on repeats, RNA-Seq mappings, transcript assembly and alignment of protein sequences was created using the REAT prediction workflow. The pipeline has four main steps: (1) the REAT transcriptome and homology Mikado models are categorised based on alignments to UniProt proteins to identify models with likely full-length CDS and which meet basic structural checks i.e., having complete but not excessively long UTRs and not exceeding a minimum CDS/cDNA ratio. A subset of gene models is then selected from the classified models and used to train the AUGUSTUS gene predictor^58^; (2) Augustus is run in both *ab initio* mode and with extrinsic evidence generated in the REAT transcriptome and homology runs (repeats, protein alignments, RNA-Seq alignments, splice junctions, categorised Mikado models). Three evidence guided AUGUSTUS predictions are created using alternative bonus scores and priority based on evidence type; (3) AUGUSTUS models, REAT transcriptome/homology models, protein and transcriptome alignments are provided to EVidenceModeler^59^ (EVM) to generate consensus gene structures; (4) EVM models are processed through Mikado to add UTR features and splice variants.

#### 5. Projection of gene models from Triticum aestivum

A reference set of hexaploid wheat gene models was derived from public gene sets (IWGSC^60^ and 10+ wheat^26^) projected onto the IWGSC RefSeq v1.0 assembly^60^; a filtered and consolidated set of models was derived with Minos, with a primary model defined for each gene. Models were scored on a combination of intrinsic gene structure characteristics, evidence support (protein and transcriptome data) and consistency in gene structure across the input gene models. The Minos primary models were classified as full-length or partial based on alignment to a filtered magnoliopsida Swiss-Prot TrEMBL database. This assignment, together with criteria for gene structure characteristics and the original confidence classification, was used to classify models into 6 categories (Platinum, Gold, Silver, Bronze, Stone and Paper), with Platinum being the highest confidence category for models assessed as full-length, with an original confidence classification of “high”, meeting structural checks for number of UTR and CDS/cDNA ratio and which were assessed as consistently annotated across the input gene sets. Reclassification resulted in 55,319 Platinum, 24,789 Gold, 11,968 Silver, 61,845 Bronze, 110,518 Stone and 115,336 Paper genes. The four highest confidence categories Platinum, Gold, Silver and Bronze were projected onto the *T. timopheevii* assembly with Liftoff v1.5.1^61^, only those models transferred fully with no loss of bases and identical exon/intron structure were retained (https://github.com/lucventurini/ei-liftover). Similarly, high confidence genes annotated in the hexaploid wheat cv. Chinese Spring RefSeq v2.1 assembly^47^ were projected onto the *T. timopheevii* genome assembly with Liftoff, and only those models transferred fully with no loss of bases and identical exon/intron structure were retained. Among these, gene models with the attribute “manually_curated” in the original Refseq v2.1 assembly were extracted as a set.

#### 6. Gene model consolidation

The final set of gene models was selected using Minos (Table 3). Minos is a pipeline that generates and utilises metrics derived from protein, transcript, and expression data sets to create a consolidated set of gene models. In this annotation, the following gene models were filtered and consolidated into a single set of gene models using Minos:

**Table 3.** Summary statistics for the final structural annotation of the *T. timopheevii* genome.

| Stat | Value |
| --- | --- |
| Number of genes | 166,325 |
| Number of Transcripts | 218,100 |
| Transcripts per gene | 1.31 |
| Number of monoexonic genes | 51,702 |
| Monoexonic transcripts | 53,192 |
| Transcript mean size cDNA (bp) | 1,658.27 |
| Transcript median size cDNA (bp) | 1412 |
| Min cDNA | 96 |
| Max cDNA | 20,589 |
| Total exons | 997,779 |
| Exons per transcript | 4.57 |
| Exon mean size (bp) | 362.47 |
| CDS mean size (bp) | 277.55 |
| Transcript mean size CDS (bp) | 1,171.61 |
| Transcript median size CDS (bp) | 957 |
| Min CDS | 0 |
| Max CDS | 20,283 |
| Intron mean size (bp) | 628.4 |
| 5'UTR mean size (bp) | 182.93 |
| 3'UTR mean size (bp) | 294.58 |

**Table 4.**
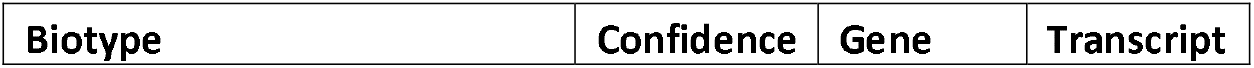
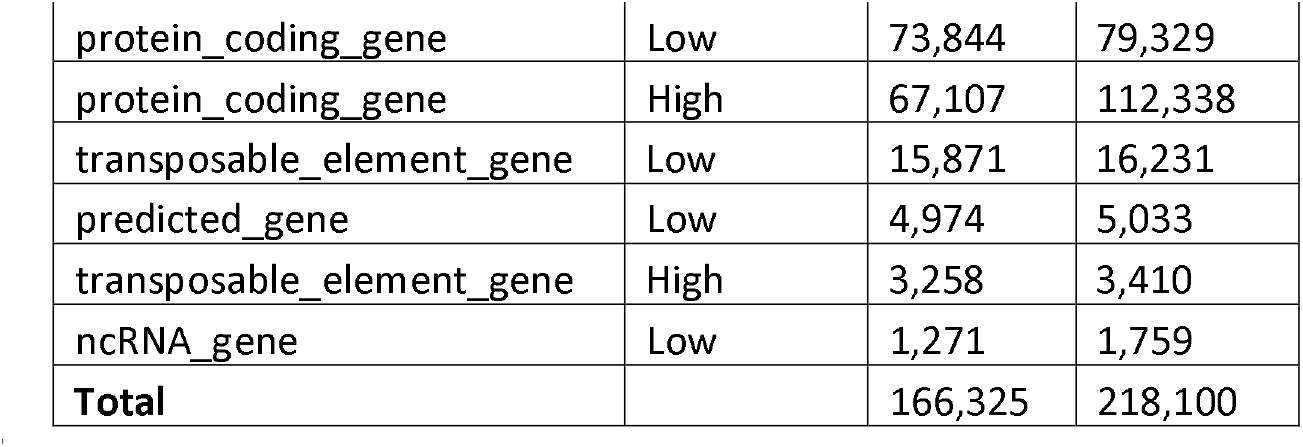
Minos classified gene models.

| Biotype | Confidence | Gene | Transcript |
| --- | --- | --- | --- |
| protein_coding_gene | Low | 73,844 | 79,329 |
| protein_coding_gene | High | 67,107 | 112,338 |
| transposable_element_gene | Low | 15,871 | 16,231 |
| predicted_gene | Low | 4,974 | 5,033 |
| transposable_element_gene | High | 3,258 | 3,410 |
| ncRNA_gene | Low | 1,271 | 1,759 |
| **Total** |  | 166,325 | 218,100 |

1. The three alternative evidence guided Augustus gene builds described earlier.
2. The gene models derived from the REAT transcriptome runs described earlier.
3. The gene models derived from the REAT homology runs described earlier.
4. The gene models derived from the REAT prediction run (AUGUSTUS and EVM-Mikado) described earlier.
5. The gene models derived from projecting public and curated *T. aestivum* gene models of varying confidence levels onto the *T. timopheevii* genome as described earlier.
6. IWGSC Refseq v2.1 models identified as “manually_curated” projected onto the *T. timopheevii* genome as described earlier.

Gene models were classified as biotypes protein_coding_gene, predicted_gene and transposable_element_gene, and assigned as high or low confidence (Table 3) based on the criteria below:

a) **High confidence (HC) protein_coding_gene:** Any protein coding gene where any of its associated gene models have a BUSCO v5.4.7^62^ protein status of Complete/Duplicated OR have diamond v0.9.36 coverage (average across query and target coverage) >= 90% against the listed Poaceae protein datasets (section 3; Supplemental File 2) or UniProt magnoliopsida proteins. Or alternatively have average blastp coverage (across query and target coverage) >= 80% against the listed protein datasets/UniProt magnoliopsida AND have transcript alignment F1 score (average across nucleotide, exon and junction F1 scores based on RNA-Seq transcript assemblies) >= 60%.
b) **Low confidence (LC) protein_coding_gene:** Any protein coding gene where all its associated transcript models do not meet the criteria to be considered as high confidence protein coding transcripts.
c) **HC transposable_element_gene:** Any protein coding gene where any of its associated gene models have coverage >= 40% against the combined interspersed repeats (see section 1).
d) **LC transposable_element_gene:** Any protein coding gene where all its associated transcript models do not meet the criteria to be considered as high confidence and assigned as a transposable_element_gene (see c).
e) **LC predicted_gene:** Any protein coding gene where all the associated transcript models do not meet the criteria to be considered as high confidence protein coding transcripts. In addition, where any of the associated gene models have average blastp coverage (across query and target coverage) < 30% against the listed protein datasets AND having a protein-coding potential score < 0.25 calculated using CPC2 0.1^63^.
f) **LC ncRNA gene:** Any gene model with no CDS features AND a protein-coding potential score < 0.3 calculated using CPC2 0.1.
g) **Discarded models:** Any models having no BUSCO protein hit AND a protein alignment score (average across nucleotide, exon and junction F1 scores based on protein alignments) <0.2 AND a transcript alignment F1 score (average across nucleotide, exon and junction F1 scores based on RNA-Seq transcript assemblies) <0.2 AND a diamond coverage (target coverage) <0.3 AND Kallisto v0.44^64^ expression score <0.2 from across RNA-Seq reads OR having short CDS <30bps. Any ncRNA genes (no CDS features) not meeting the ncRNA gene requirements (f) were also excluded.

Gene model distribution across the pseudomolecules and unplaced scaffolds is shown in Table 2 and gene density of 164,617 protein coding genes across the *T. timopheevii* genome is shown in Fig. 2b.

**Figure 2.**
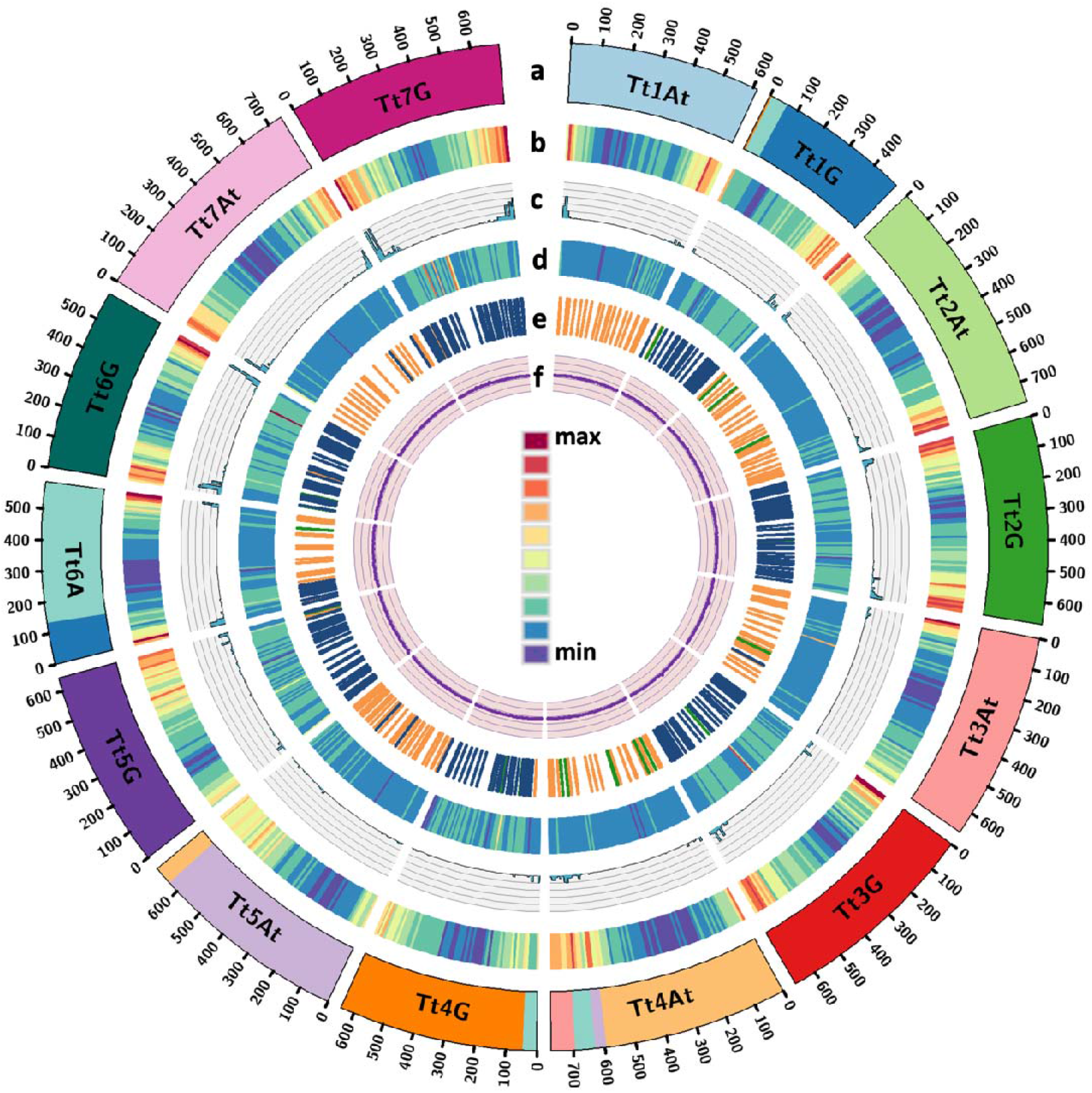
Circos plot^71^ of features of the chromosome-scale assembly of *T. timopheevii* showing (a) major translocations with the *T. timopheevii* genome as observed through collinearity analysis against *T. turgidum*, (b) gene density (of all gene models), (c) NLR density (max count 87), (d) DNA methylation (5mC modification) density, (e) distribution of KASP markers based on SNPs with bread wheat cv. Chinese Spring^29^ and (f) GC content. Tt in chromosome name represents *T. timopheevii*.

#### 7. Functional annotation

All proteins were annotated using AHRD v.3.3.3 (available at https://github.com/groupschoof/AHRD/blob/master/README.textile). Sequences were compared against the reference proteins (Arabidopsis thaliana TAIR10, TAIR10_pep_20101214_updated.fasta.gz - https://www.araport.org) and the UniProt viridiplantae sequences^65^ (data download 06-May-2023), both Swiss-Prot and TrEMBL datasets using blastp v2.6.0 with an e-value of 1e-5. InterproScan v5.22.61^66^ results were also provided to AHRD. The standard AHRD example configuration file path test/resources/ahrd_example_input_go_prediction.yml, distributed with the AHRD tool, was adapted apart from the location of input and output files. The GOA mapping from UniProt (ftp://ftp.ebi.ac.uk/pub/databases/GO/goa/UNIPROT/goa_uniprot_all.gaf.gz) was included as parameter ‘gene_ontology_result’. The interpro database (ftp://ftp.ebi.ac.uk/pub/databases/interpro/61.0/interpro.xml.gz) was included as parameter ‘interpro_database’. The parameter ‘prefer_reference_with_go_annos’ was changed to ‘false’ and the blast database specific weights used were:

blast_dbs:

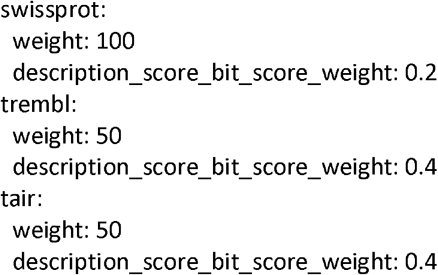

Since *T. timopheevii* is known as an important source for genetic variation for resistance against major diseases of wheat as described above and as the majority of cloned disease-resistance genes encode nucleotide-binding leucine-rich repeats (NLRs)^67,68^, we analysed the genomic distribution of all gene models annotated as NB-ARC domain-containing/disease resistance proteins in the genome assembly (Fig. 2c).

### Generation of PacBio DNA methylation profile

Methylation in CpG context was inferred with ccsmeth v0.3.2^69^, using the kinetics data from PacBio CCS subreads obtained during HMW DNA sequencing. The methylation prediction for CCS reads were called using the model “model_ccsmeth_5mCpG_call_mods_attbigru2s_b21.v2.ckpt”. The reads with the MM⍰+⍰ML tags were aligned to the pseudomolecules in the *T. timopheevii* assembly using BWA v0.7.17^70^. The methylation frequency was calculated at genome level with the modbam files and the aggregate mode of ccsmeth with the model “model_ccsmeth_5mCpG_aggregate_attbigru_b11.v2p.ckpt”. The genomic distribution of 5mC modifications across *T. timopheevii* (Fig. 2d) shows that G genome chromosomes have more methylation with an average of ∼401.8 Kb methylated bases per 10 Mb bin as compared to the A^t^ genome chromosomes with an average of ∼385.5 Kb per 10 Mb bin.

### Comparative genome analysis

Synteny and collinearity analysis of the *T. timopheevii* gene set against the reference gene set of wild emmer wheat *T. turgidum* accession Zavitan WEWSeq v1.0^72,73^ (available from Ensembl^74^ plants) was computed using MCScanX^75^ with defaults parameters and results viewed using SynVisio^76,77^ (https://synvisio.github.io/) and shown in Fig. 3a-b. A and G genome chromosomes of *T. timopheevii* maintain synteny with the A and B genome chromosomes of tetraploid wheat albeit some inversions, deletions and translocations shown in red Fig. 3a. Analysis of large chromosome translocations within the *T. timopheevii* genome confirmed previous reports^7-9^ of 5 translocation events including T4A^t^L/5A^t^L (1), T6A^t^S/1GS/4GS (2-4) and T4A^t^L/3A^t^L (5). Fig. 3b shows the composition of the chromosomes involved (or suspected to be) in the translocation events as compared to the composition of homoeologous chromosomes in tetraploid wheat (also depicted in Fig. 2a). It shows that Chr4GS had retained a part of Chr6A^t^S during the fourth reciprocal translocation event between T4A^t^L/4GS^8^ unlike previous reports that indicated that all of Chr6A^t^S was translocated to Chr4A^t^L. It was also confirmed that unlike tetraploid (and hexaploid) wheat there is no inversion of Chr4A^t^L (also shown in Fig. 3c) and no reciprocal translocation between Chr7G and Chr4A^t^L^78,79^ indicating that although the T4AL/5AL was inherited from *T. urartu*^8,80^, the following inversion of Chr4AL and translocation with Chr7B were specific to the tetraploid and hexaploid wheat species.

**Figure 3.**
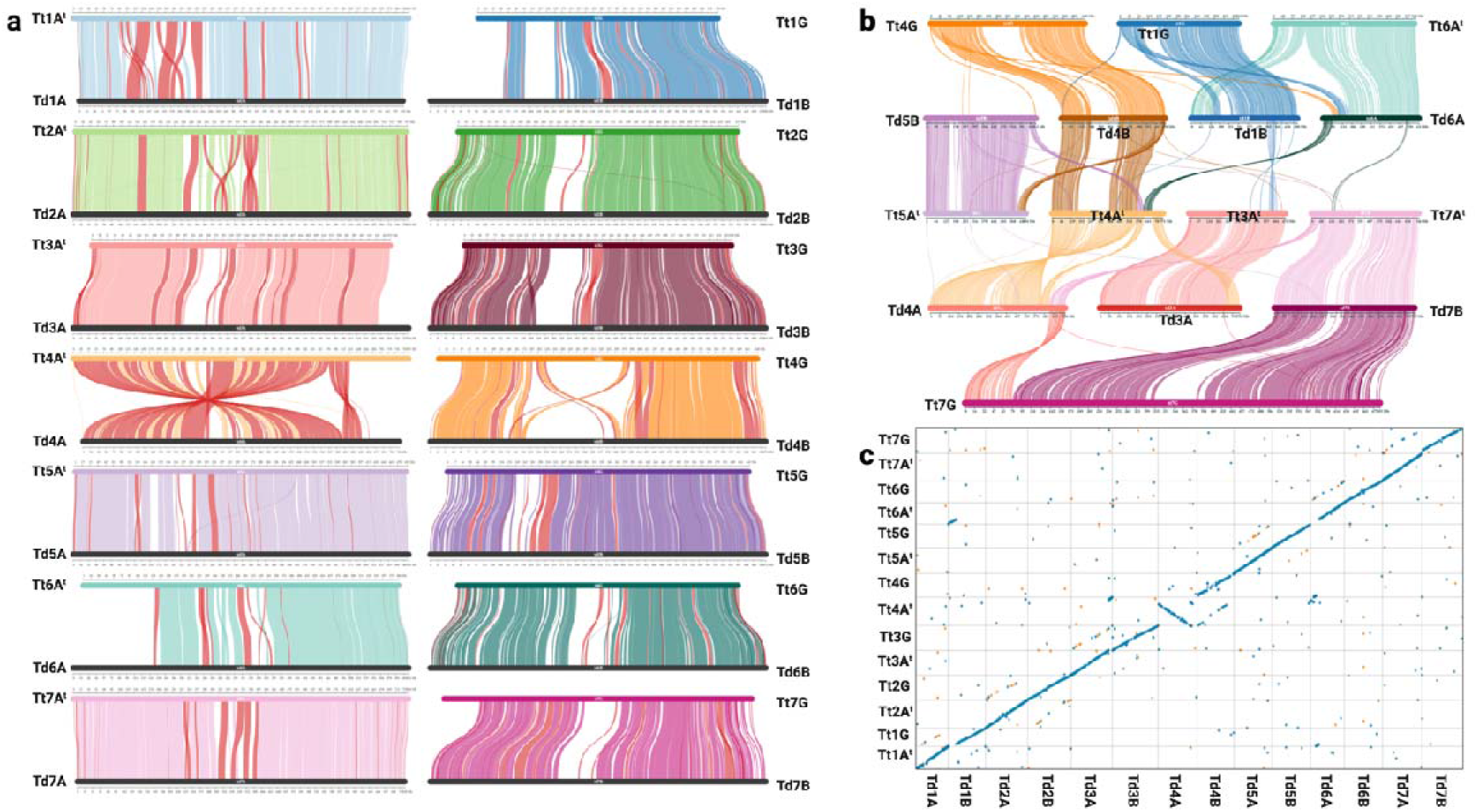
Comparative analysis of *T. timopheevii* (Tt) and *T. turgidum* (Td) genomes. (a) SynVisio plots showing synteny and collinearity between the two genomes with rearrangements in red, (b) SynVisio plots showing major translocations within the *T. timopheevii* chromosomes as compared to tetraploid wheat, and (c) Dotplot comparison of the sequence alignments between the chromosomes of the two genomes.

Dotplot comparison of sequence alignments (as described earlier) between the *T. timopheevii* pseudomolecules and the reference genome of *T. turgidum* accession Zavitan^72,73^ WEWSeq v1.0 also confirmed the synteny, collinearity and translocations (Fig. 3c) as observed by comparing the gene sets between these two species (Fig. 3a-b).

### Phylogeny analysis

Orthofinder^81^ (https://github.com/davidemms/OrthoFinder) was used to locate orthologous genes between *T. timopheevii* (Tt)and other Aegilops and wheat annotations using protein coding genes (HC + LC). We used the S genome annotations from three *Aegilops* species^82^: Ae. longissima, Ae. speltoides and *Ae. sharonensis*, the A genome annotation of *T. urartu*, the D genome annotation of Ae.⍰tauschii, the AB annotation of wild emmer wheat (WEW) accession Zavitan^72^ and durum wheat (DW) cv. Svevo^83^ and the ABD annotation of hexaploid bread wheat (BW) cv. Chinese Spring^60^ (available from Ensembl plants). The polyploid annotations (*T. timopheevii*, WEW, DW and BW) were split into the subgenomes, and each was handled separately. The species tree (Fig. 4) was viewed using the ETE Toolkit tree viewer^84^ (available at http://etetoolkit.org/treeview/) and confirms that the G genome of *T. timopheevii* is more closely related to the S genome of *Ae. speltoides* than the B genomes of tetraploid and hexaploid wheats.

**Figure 4.**
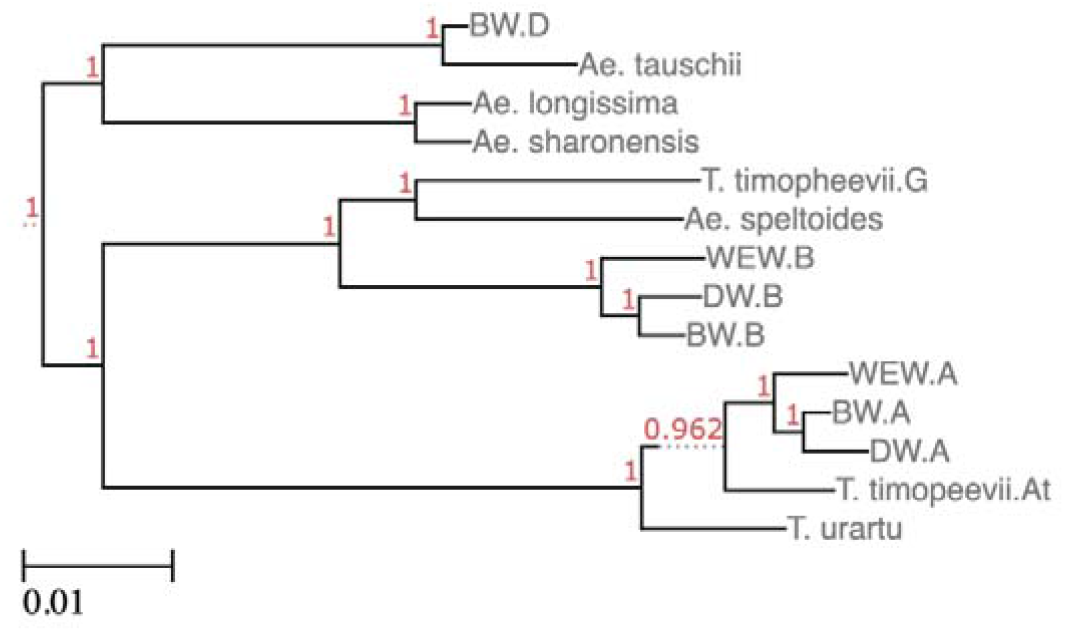
Phylogenetic tree based on orthofinder analysis of all protein coding genes. Branch values in red correspond to orthofinder support values. BW, bread wheat cv. Chinese Spring; DW durum wheat cv. Svevo; WEW, wild emmer wheat accession Zavitan.

### Genome visualisation

A genome browser for the assembly of *T. timopheevii* generated in this study is currently being hosted at GrainGenes^85^ (https://wheat.pw.usda.gov/jb?data=/ggds/whe-timopheevii) with tracks for annotated gene models and repeats and BLAST functionality available at https://wheat.pw.usda.gov/blast/.

### Data Records

The raw sequence files for the HiFi, Hi-C, RNA-Seq and IsoSeq reads were deposited in the European Nucleotide Archive (ENA) under accession number <u>PRJEB71660</u>. The final chromosome-scale assembly consisting of the nuclear and organelle genomes was deposited at ENA under the accession number GCA_963921465.2.

The genome assemblies, gene model and repeat annotations, methylation profile and Hi-C contact map are also available at on DRYAD Digital Repository^86^ (https://doi.org/10.5061/dryad.mpg4f4r6p).

## Technical Validation

### Assessment of genome assembly and annotation

The final curated assembly was assessed by mapping the trimmed Hi-C reads to the post-curated assembly (as described above for scaffolding) and generating a final Hi-C contact map using PretextMap v0.1.9 and viewed using PretextView v0.2.5. It showed a dense dark blue pattern along the diagonal revealing no potential mis-assemblies (Fig. 5). The anti-diagonals in the Hi-C contact matrix were expected and have been reported for other relatively large plant genomes such as those from the Triticeae tribe^87,88^ as they correspond to the typical Rabl configuration of Triticeae chromosomes^89,90^.

**Figure 5.**
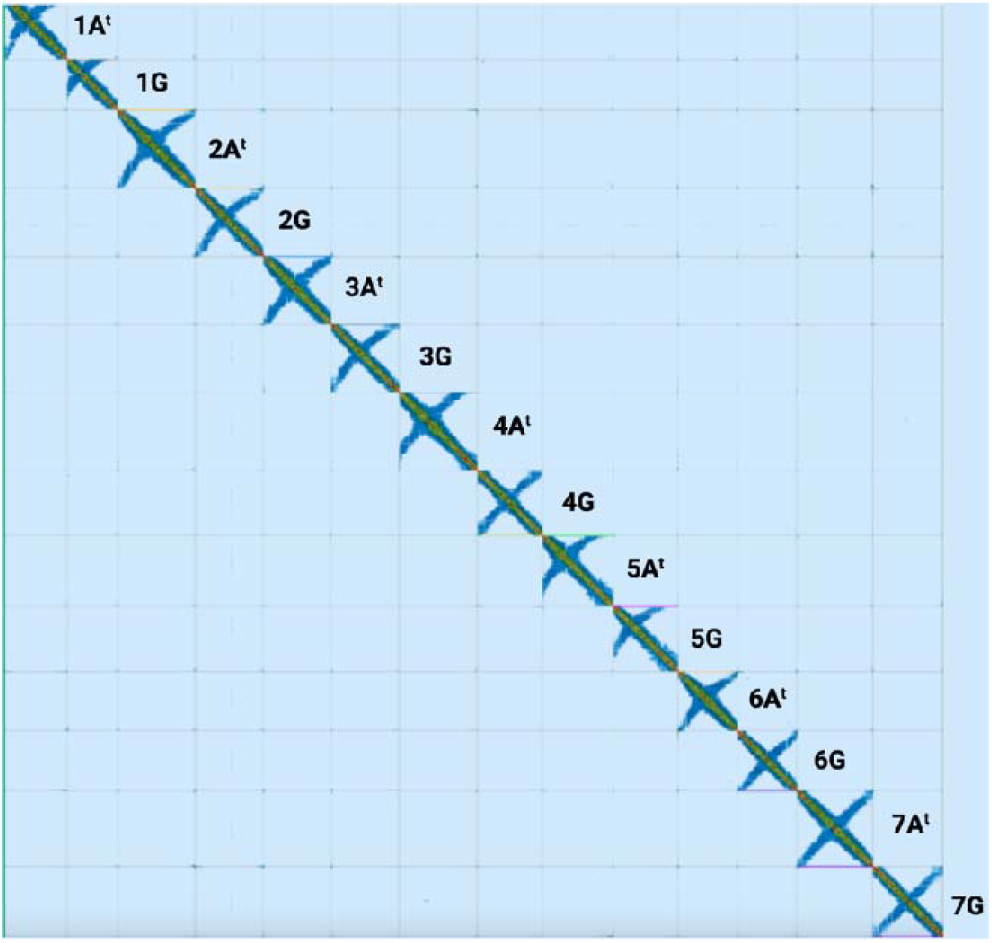
Contact map after the integration of the Hi-C data and manual correction using PretextView.

The BUSCO v5.3.2^42^ (-l embryophyta_odb10) score of 98.9% (0.6% fragmented and 0.5% missing BUSCOs; Table S8) at the genome level indicates a high completeness of the *T. timopheevii* assembly. The quality of the *T. timopheevii* assembly was assessed with Merqury^91^ based on the PacBio HiFi reads using 31-mers. The QV (consensus quality value) and k-mer completeness scores were 65.5 and 97.8%, respectively.

Completeness of the gene model prediction was also evaluated using BUSCO and produced a score of 99.7% (0.1% fragmented and 0.2% missing BUSCOs; Table S8). The number of HC gene models (70,365) is in the range of a tetraploid Triticeae species (34,000–43,000 high-confidence gene models per haploid genome)^92^.

Of the total 14 chromosomes, we found telomeric repeats on both ends for 5 chromosomes (1A^t^, 2G, 3A^t^, 6A^t^, and 7G) and on one end for 7 chromosomes (1GL, 2A^t^S, 3GL, 4GS, 5GL, 6GL and 7A^t^L).

## Code availability

All software and pipelines were executed according to the manual and protocol of published tools. No custom code was generated for these analyses.

## Acknowledgements

This work was supported by the Biotechnology and Biological Sciences Research Council [grant number BB/P016855/1] as part of the Developing Future Wheat (DFW) programme. Part of this work was also delivered via Transformative Genomics the BBSRC funded National Bioscience Research Infrastructure (BBS/E/ER/23NB0006) at Earlham Institute by members of the Genomics Pipelines and Core Bioinformatics Groups. EY and TS were supported by the US. Department of Agriculture, Agricultural Research Service, Project No. 2030–21000-056-00D.

## Author contributions

SuG, JK and IK designed the study and obtained funding for it. CY, DuS and SA carried out plant maintenance and nucleic acid extraction. SuG, MW and LY generated the genome assembly. SuG and JC carried out manual curation of the assembly. SrG and DaS carried out the genome annotation. EY and TS generated the genome browser. SuG wrote the initial manuscript. All authors have read and approved the final manuscript.

## References

1. Dvořák, J., Terlizzi, P. d., Zhang, H.-B., Resta, P. The evolution of polyploid wheats: identification of the A genome donor species. Genome 36, 21–31 (1993).

2. Dvorak, J., Zhang, H.-B. Variation in repeated nucleotide sequences sheds light on the phylogeny of the wheat B and G genomes. Proceedings of the National Academy of Sciences 87, 9640–9644 (1990).

3. Ahmed, H. I., et al. Einkorn genomics sheds light on history of the oldest domesticated wheat. Nature 620, 830–838 (2023).

4. Rodriguez, S., Maestra, B., Perera, E., Diez, M., Naranjo, T. Pairing affinities of the B-and G-genome chromosomes of polyploid wheats with those of Aegilops speltoides. Genome 43, 814–819 (2000).

5. Li, L. F., et al. Genome sequences of five Sitopsis species of Aegilops and the origin of polyploid wheat B subgenome. Molecular plant 15, 488–503 (2022).

6. Dvořák, J. Triticum Species (Wheat). Encyclopedia of Genetics, 2060–2068 (2001).

7. Jiang, J., Gill, B. S. Different species-specific chromosome translocations inTriticum timopheevii and T. turgidum support the diphyletic origin of polyploid wheats. Chromosome Research 2, 59–64 (1994).

8. Maestra, B., Naranjo, T. Structural chromosome differentiation between Triticum timopheevii and T. turgidum and T. aestivum. Theoretical and Applied Genetics 98, 744–750 (1999).

9. Rodriguez, S., Perera, E., Maestra, B., Díez, M., Naranjo, T. Chromosome structure of Triticum timopheevii relative to T. turgidum. Genome 43, 923–930 (2000).

10. Devi, U., et al. Development and characterisation of interspecific hybrid lines with genome-wide introgressions from Triticum timopheevii in a hexaploid wheat background. BMC Plant Biol 19, 183 (2019).

11. Brown-Guedira, G. L., Singh, S., Fritz, A. K. Performance and Mapping of Leaf Rust Resistance Transferred to Wheat from Triticum timopheevii subsp. armeniacum. Phytopathology 93, 784–789 (2003).

12. Singh, A. K., et al. Genetics and mapping of a new leaf rust resistance gene in Triticum aestivum L. × Triticum timopheevii Zhuk. derivative ‘Selection G12’. J Genet 96, 291–297 (2017).

13. Leonova, I. N., et al. Microsatellite mapping of a leaf rust resistance gene transferred to common wheat from Triticum timopheevii. Cereal Research Communications 38, 211–219 (2010).

14. McIntosh, R., Gyarfas, J. Triticum timopheevii as a source of resistance to wheat stem rust. Zeitschrift fur Pflanzenzuchtung 66, 240–248 (1971).

15. Wu, S., Pumphrey, M., Bai, G. Molecular Mapping of Stem-Rust-Resistance Gene Sr40 in Wheat. Crop Science 49, 1681–1686 (2009).

16. Allard, R., Shands, R. Inheritance of resistance to stem rust and powdery mildew in cytologically stable spring wheats derived from Triticum timopheevii. Phytopathology 44, 266–274 (1954).

17. Perugini, L. D., Murphy, J. P., Marshall, D., Brown-Guedira, G. Pm37, a new broadly effective powdery mildew resistance gene from Triticum timopheevii. Theoretical and Applied Genetics 116, 417–425 (2008).

18. Qin, B., et al. Collinearity-based marker mining for the fine mapping of Pm6, a powdery mildew resistance gene in wheat. Theoretical and Applied Genetics 123, 207–218 (2011).

19. Steed, A., et al. Identification of Fusarium Head Blight Resistance in Triticum timopheevii Accessions and Characterization of Wheat-T. timopheevii Introgression Lines for Enhanced Resistance. Frontiers in Plant Science 13, (2022).

20. Malihipour, A., Gilbert, J., Fedak, G., Brûlé-Babel, A., Cao, W. Characterization of agronomic traits in a population of wheat derived from Triticum timopheevii and their association with Fusarium head blight. European Journal of Plant Pathology 144, 31–43 (2016).

21. Brown-Guedira, G., et al. Evaluation of a collection of wild timopheevi wheat for resistance to disease and arthropod pests. Plant disease 80, 928–933 (1996).

22. Badridze, G., Weidner, A., Asch, F., Börner, A. Variation in salt tolerance within a Georgian wheat germplasm collection. Genetic resources and crop evolution 56, 1125–1130 (2009).

23. Yudina, R., Leonova, I., Salina, E., Khlestkina, E. Change in salt tolerance of bread wheat as a result of the introgression of the genetic material of Aegilops speltoides and Triticum timopheevii. Russian Journal of Genetics: Applied Research 6, 244–248 (2016).

24. Lehmensiek, A., Bovill, W., Banks, P., Sutherland, M. Molecular characterization of a Triticum timopheevii introgression in a Wentworth/Lang population. (2008).

25. Hu, X., et al. Zn and Fe concentration variations of grain and flag leaf and the relationship with NAM-G1 gene in Triticum timopheevii (Zhuk.) Zhuk. ssp. *timopheevii*. Cereal Research Communications 45, 421–431 (2017).

26. Walkowiak, S., et al. Multiple wheat genomes reveal global variation in modern breeding. Nature 588, 277–283 (2020).

27. Keilwagen, J., et al. Detecting major introgressions in wheat and their putative origins using coverage analysis. Scientific Reports 12, 1908 (2022).

28. Keilwagen, J., et al. Finding needles in a haystack: identification of inter-specific introgressions in wheat genebank collections using low-coverage sequencing data. Frontiers in Plant Science 14, (2023).

29. King, J., et al. Introgression of the Triticum timopheevii Genome Into Wheat Detected by Chromosome-Specific Kompetitive Allele Specific PCR Markers. Frontiers in Plant Science 13, (2022).

30. Grewal, S., et al. Rapid identification of homozygosity and site of wild relative introgressions in wheat through chromosome-specific KASP genotyping assays. Plant Biotechnol J 18, 743–755 (2020).

31. Belton, J. M., et al. Hi-C: a comprehensive technique to capture the conformation of genomes. Methods 58, 268–276 (2012).

32. Wenger, A. M., et al. Accurate circular consensus long-read sequencing improves variant detection and assembly of a human genome. Nature Biotechnology 37, 1155–1162 (2019).

33. Driguez, P., et al. LeafGo: Leaf to Genome, a quick workflow to produce high-quality de novo plant genomes using long-read sequencing technology. Genome biology 22, 256 (2021).

34. Dong, L., et al. Single-molecule real-time transcript sequencing facilitates common wheat genome annotation and grain transcriptome research. BMC Genomics 16, 1039 (2015).

35. Martin, M. Cutadapt removes adapter sequences from high-throughput sequencing reads. 2011 17, 3 (2011).

36. Bolger, A. M., Lohse, M., Usadel, B. Trimmomatic: a flexible trimmer for Illumina sequence data. Bioinformatics 30, 2114–2120 (2014).

37. Marçais, G., Kingsford, C. A fast, lock-free approach for efficient parallel counting of occurrences of k-mers. Bioinformatics 27, 764–770 (2011).

38. Wang, H., et al. Estimation of genome size using k-mer frequencies from corrected long reads. arXiv:200311817 [q-bioGN], (2020).

39. Ranallo-Benavidez, T. R., Jaron, K. S., Schatz, M. C. GenomeScope 2.0 and Smudgeplot for reference-free profiling of polyploid genomes. Nature Communications 11, 1432 (2020).

40. Cheng, H., Concepcion, G. T., Feng, X., Zhang, H., Li, H. Haplotype-resolved de novo assembly using phased assembly graphs with hifiasm. Nature Methods 18, 170–175 (2021).

41. Formenti, G., et al. Gfastats: conversion, evaluation and manipulation of genome sequences using assembly graphs. Bioinformatics 38, 4214–4216 (2022).

42. Simão, F. A., Waterhouse, R. M., Ioannidis, P., Kriventseva, E. V., Zdobnov, E. M. BUSCO: assessing genome assembly and annotation completeness with single-copy orthologs. Bioinformatics 31, 3210–3212 (2015).

43. Laetsch, D., Blaxter, M. BlobTools: Interrogation of genome assemblies. F1000Research 6, (2017).

44. Korbel, J. O., Lee, C. Genome assembly and haplotyping with Hi-C. Nature Biotechnology 31, 1099–1101 (2013).

45. Ghurye, J., et al. Integrating Hi-C links with assembly graphs for chromosome-scale assembly. PLOS Computational Biology 15, e1007273 (2019).

46. Howe, K., et al. Significantly improving the quality of genome assemblies through curation. GigaScience 10, (2021).

47. Zhu, T., et al. Optical maps refine the bread wheat Triticum aestivum cv. Chinese Spring genome assembly. The Plant Journal 107, 303–314 (2021).

48. Kurtz, S., et al. Versatile and open software for comparing large genomes. Genome biology 5, R12 (2004).

49. Venturini, L., Caim, S., Kaithakottil, G. G., Mapleson, D. L., Swarbreck, D. Leveraging multiple transcriptome assembly methods for improved gene structure annotation. GigaScience 7, (2018).

50. Boden, S. A., et al. Updated guidelines for gene nomenclature in wheat. Theoretical and Applied Genetics 136, 72 (2023).

51. Kim, D., Paggi, J. M., Park, C., Bennett, C., Salzberg, S. L. Graph-based genome alignment and genotyping with HISAT2 and HISAT-genotype. Nature Biotechnology 37, 907–915 (2019).

52. Li, H. Minimap2: pairwise alignment for nucleotide sequences. Bioinformatics 34, 3094–3100 (2018).

53. Mapleson, D., Venturini, L., Kaithakottil, G., Swarbreck, D. Efficient and accurate detection of splice junctions from RNA-seq with Portcullis. GigaScience 7, (2018).

54. Kovaka, S., et al. Transcriptome assembly from long-read RNA-seq alignments with StringTie2. Genome biology 20, 278 (2019).

55. Shao, M., Kingsford, C. Accurate assembly of transcripts through phase-preserving graph decomposition. Nature Biotechnology 35, 1167–1169 (2017).

56. Gotoh, O. A space-efficient and accurate method for mapping and aligning cDNA sequences onto genomic sequence. Nucleic Acids Research 36, 2630–2638 (2008).

57. Li, H. Protein-to-genome alignment with miniprot. Bioinformatics 39, (2023).

58. Stanke, M., Morgenstern, B. AUGUSTUS: a web server for gene prediction in eukaryotes that allows user-defined constraints. Nucleic Acids Research 33, W465–W467 (2005).

59. Haas, B. J., et al. Automated eukaryotic gene structure annotation using EVidenceModeler and the Program to Assemble Spliced Alignments. Genome biology 9, R7 (2008).

60. IWGSC, et al. Shifting the limits in wheat research and breeding using a fully annotated reference genome. Science 361, (2018).

61. Shumate, A., Salzberg, S. L. Liftoff: accurate mapping of gene annotations. Bioinformatics 37, 1639–1643 (2021).

62. Seppey, M., Manni, M., Zdobnov, E. M. in Gene Prediction: Methods and Protocols (ed. Kollmar M) BUSCO: Assessing Genome Assembly and Annotation Completeness (Springer New York, 2019).

63. Kong, L., et al. CPC: assess the protein-coding potential of transcripts using sequence features and support vector machine. Nucleic Acids Research 35, W345–W349 (2007).

64. Bray, N. L., Pimentel, H., Melsted, P., Pachter, L. Near-optimal probabilistic RNA-seq quantification. Nature Biotechnology 34, 525–527 (2016).

65. Consortium, U. UniProt: a hub for protein information. Nucleic Acids Res 43, D204–212 (2015).

66. Jones, P., et al. InterProScan 5: genome-scale protein function classification. Bioinformatics 30, 1236–1240 (2014).

67. Kourelis, J., Van Der Hoorn, R. A. Defended to the nines: 25 years of resistance gene cloning identifies nine mechanisms for R protein function. The Plant cell 30, 285–299 (2018).

68. Chen, R., Gajendiran, K., Wulff, B. B. H. R we there yet? Advances in cloning resistance genes for engineering immunity in crop plants. Current opinion in plant biology 77, 102489 (2024).

69. Ni, P., et al. DNA 5-methylcytosine detection and methylation phasing using PacBio circular consensus sequencing. Nature communications 14, 4054 (2023).

70. Li, H., Durbin, R. Fast and accurate short read alignment with Burrows–Wheeler transform. bioinformatics 25, 1754–1760 (2009).

71. Krzywinski, M., et al. Circos: An information aesthetic for comparative genomics. Genome Res 19, 1639–1645 (2009).

72. Avni, R., et al. Wild emmer genome architecture and diversity elucidate wheat evolution and domestication. Science 357, 93–97 (2017).

73. Zhu, T., et al. Improved Genome Sequence of Wild Emmer Wheat Zavitan with the Aid of Optical Maps. G3 (Bethesda) 9, 619–624 (2019).

74. Martin, F. J., et al. Ensembl 2023. Nucleic Acids Research 51, D933–D941 (2022).

75. Wang, Y., et al. MCScanX: a toolkit for detection and evolutionary analysis of gene synteny and collinearity. Nucleic acids research 40, e49–e49 (2012).

76. Bandi, V., Gutwin, C. Interactive Exploration of Genomic Conservation. In Proceedings of 46th Graphics Interface Conference (Canadian Human-Computer Communications Society, 2020).

77. Bandi, V., et al. in Plant Bioinformatics: Methods and Protocols (ed. Edwards D) Visualization Tools for Genomic Conservation (Springer US, 2022).

78. Devos, K. M., Dubcovsky, J., Dvorak, J., Chinoy, C. N., Gale, M. D. Structural evolution of wheat chromosomes 4A, 5A, and 7B and its impact on recombination. Theor Appl Genet 91, 282–288 (1995).

79. Dvorak, J., et al. Reassessment of the evolution of wheat chromosomes 4A, 5A, and 7B. Theor Appl Genet 131, 2451–2462 (2018).

80. King, I. P., et al. Detection of interchromosomal translocations within the Triticeae by RFLP analysis. Genome 37, 882–887 (1994).

81. Emms, D., Kelly, S. OrthoFinder: phylogenetic orthology inference for comparative genomics. bioRxiv 466201, 2019.

82. Avni, R., et al. Genome sequences of three Aegilops species of the section Sitopsis reveal phylogenetic relationships and provide resources for wheat improvement. The Plant Journal 110, 179–192 (2022).

83. Maccaferri, M., et al. Durum wheat genome highlights past domestication signatures and future improvement targets. Nature Genetics 51, 885–895 (2019).

84. Huerta-Cepas, J., Serra, F., Bork, P. ETE 3: Reconstruction, Analysis, and Visualization of Phylogenomic Data. Molecular Biology and Evolution 33, 1635–1638 (2016).

85. Yao, E., et al. GrainGenes: a data-rich repository for small grains genetics and genomics. Database 2022, (2022).

86. Grewal, S., et al. 2024. Triticum timopheevii genome assembly files [Dataset]. Dryad. 10.5061/dryad.mpg4f4r6p

87. Dong, P., et al. 3D chromatin architecture of large plant genomes determined by local A/B compartments. Molecular plant 10, 1497–1509 (2017).

88. Mascher, M., et al. A chromosome conformation capture ordered sequence of the barley genome. Nature 544, 427–433 (2017).

89. Anamthawat-Jónsson, K., Heslop-Harrison, J. Centromeres, telomeres and chromatin in the interphase nucleus of cereals. Caryologia 43, 205–213 (1990).

90. Cowan, C. R., Carlton, P. M., Cande, W. Z. The polar arrangement of telomeres in interphase and meiosis. Rabl organization and the bouquet. Plant Physiology 125, 532–538 (2001).

91. Rhie, A., Walenz, B. P., Koren, S., Phillippy, A. M. Merqury: reference-free quality, completeness, and phasing assessment for genome assemblies. Genome biology 21, 1–27 (2020).

92. Poretti, M., Praz, C. R., Sotiropoulos, A. G., Wicker, T. A survey of lineage-specific genes in Triticeae reveals de novo gene evolution from genomic raw material. Plant Direct 7, e484 (2023).

