## Supplemental Tables S1-S8 for "Chromosome-scale genome assembly of bread wheat’s wild relative *Triticum timopheevii*"

**Table S1.** Statistics of HiFi reads.

| **Sample** | **Cell** | **HiFi reads bases (bp)** | **Total bases (Gb)** | **HiFi reads number** | **Average HiFi reads length** | **N50** |
| --- | --- | --- | --- | --- | --- | --- |
| S95 | S95_m64165_220422_130103 | 26,589,037,091 | 199.12 | 1,842,033 | 14,434 | 16,850 |
| S95 | S95_m64165_220425_132611 | 10,060,161,095 |  | 640,383 | 15,709 | 16,775 |
| S95 | S95_m64164_220605_052635 | 22,093,742,922 |  | 1,684,360 | 13,116 | 15,031 |
| S95 | S95_m64164_220610_124716 | 21,036,575,173 |  | 1,478,325 | 14,230 | 15,571 |
| S95 | S95_m64164_220611_190014 | 28,784,821,607 |  | 1,886,131 | 15,261 | 15,752 |
| S95 | S95_m64164_220613_011317 | 32,085,349,484 |  | 2,123,969 | 15,106 | 15,590 |
| S95 | S95_m64164_220615_140016 | 27,316,613,580 |  | 1,919,234 | 14,233 | 15,115 |
| S95 | S95_m64165_220615_155227 | 31,158,294,078 |  | 2,066,458 | 15,078 | 15,482 |
| P95 | P95_m64165_221012_052633 | 35,716,109,179 | 67.53 | 1,962,144 | 18,202 | 18,092 |
| P95 | P95_m64267e_221023_073301 | 31,810,292,249 |  | 1,782,735 | 17,843 | 17,713 |
|  | **Total** | 266,650,996,458 | 267 | 17,385,772 |  |  |

**Table S2.** Statistics of HiC reads.

| **Sample** | **Library_Flowcell_Lane** | **Raw reads (bp)** | **Raw data (Gb)** | **Effective(%)** | **Error(%)** | **Q20(%)** | **Q30(%)** | **GC(%)** |
| --- | --- | --- | --- | --- | --- | --- | --- | --- |
| sp22161_1 | EKDL220006552-1a_HKG5JDSX3_L4 | 177,231,714 | 214.1 | 100 | 0.03 | 96.32 | 90.45 | 45.66 |
| sp22161_1 | EKDL220006552-1a_HKHCTDSX3_L1 | 1,250,355,254 |  | 99.35 | 0.03 | 94.71 | 88.33 | 46.04 |
| sp22161_2 | EKDL220006549-1a_HKHCTDSX3_L1 | 1,350,683,438 | 202.6 | 99.35 | 0.03 | 94.95 | 88.94 | 46.69 |
| sp22161_3 | EKDL220006550-1a_HKG5JDSX3_L4 | 137,595,592 | 212.1 | 99.99 | 0.03 | 96.87 | 92.09 | 46.45 |
| sp22161_3 | EKDL220006550-1a_HKHCTDSX3_L1 | 1,276,257,308 |  | 99.32 | 0.03 | 95.06 | 89.42 | 46.62 |
| sp22161_4 | EKDL220006551-1a_HKHCTDSX3_L1 | 1,144,706,830 | 213.4 | 99.37 | 0.03 | 94.37 | 87.99 | 46.5 |
| sp22161_4 | EKDL220006551-1a_HKG2JDSX3_L3 | 277,860,482 |  | 100 | 0.03 | 95.93 | 89.76 | 46.15 |
|  | **Total** | 5,614,690,618 | 842.2 |  |  |  |  |  |

**Table S3.** Statistics of mRNA sequencing.

| **Sample*** | **Library_Flowcell_Lane** | **Raw reads (bp)** | **Raw data (Gb)** | **Effective(%)** | **Error(%)** | **Q20(%)** | **Q30(%)** | **GC(%)** |
| --- | --- | --- | --- | --- | --- | --- | --- | --- |
| Tim_Grn | EKRN230001853-1A_HMTFTDSX5_L2 | 29,753,674 | 61.5 | 96.63 | 0.03 | 96.03 | 90.63 | 52.03 |
| Tim_Grn | EKRN230001853-1A_HMTHFDSX5_L2 | 380,224,686 |  | 97.55 | 0.03 | 97.17 | 93.17 | 52.72 |
| Tim_RT | EKRN230001854-1A_HMTHFDSX5_L2 | 528,260,132 | 79.2 | 98.38 | 0.02 | 98.08 | 94.75 | 55.62 |
| Tim_Ypl_am | EKRN230001855-1A_HMTHFDSX5_L2 | 489,608,798 | 73.4 | 98.56 | 0.02 | 98.12 | 94.84 | 56.85 |
| Tim_Ypl_pm | EKRN230001856-1A_HMTHFDSX5_L2 | 352,306,000 | 62.5 | 98.78 | 0.03 | 97.78 | 93.98 | 56.07 |
| Tim_Ypl_pm | EKRN230001856-1A_HMTFTDSX5_L2 | 64,037,984 |  | 98.27 | 0.03 | 96.46 | 90.91 | 55.23 |
| Tim_Spk | EKRN230001857-1A_HMTHFDSX5_L2 | 446,443,460 | 67 | 98.57 | 0.02 | 98.03 | 94.6 | 55.08 |
| Tim_Flg | EKRN230001852-1A_HMVJWDSX5_L2 | 406,828,476 | 61 | 97.87 | 0.03 | 97.37 | 93.15 | 56.65 |
|  | **Total** | 2,697,463,210 | 405 |  |  |  |  |  |

| ***Sample_Code** | **Sample_Description** |
| --- | --- |
| Tim_Flg | Timopheevii Flag Leaf |
| Tim_Grn | Timopheevii Grains |
| Tim_RT | Timopheevii Roots |
| Tim_Ypl_am | Timopheevii seedlings dawn |
| Tim_Ypl_pm | Timopheevii seedlings dusk |
| Tim_Spk | Timopheevii Spike |

**Table S4.** Statistics of Iso-Seq sequencing (a) and initial analysis (b) using the PacBio Iso-Seq pipeline.

(a)

| **Sample** | **Cell** | **HiFi reads bases (bp)** | **Total bases(Gb)** | **HiFi reads number** | **Average HiFi reads length** | **N50** |
| --- | --- | --- | --- | --- | --- | --- |
| Tim_pool | Tim_pool_m64165_230216_134625 | 4,468,440,060 | 4.47 | 2,179,791 | 2,049 | 2,216 |

(b)

| **CCS Analysis Read Classification** | **Values** |
| --- | --- |
| Reads | 2,179,791 |
| Reads with 5' and 3' Primers | 2,013,464 |
| Non-Concatamer Reads with 5' and 3' Primers | 2,010,455 |
| Non-Concatamer Reads with 5' and 3' Primers and Poly-A Tail | 2,008,983 |
| Mean Length of Full-Length Non-Concatamer Reads | 1,974 |
| Unique Primers | 1 |
| Mean Reads per Primer | 2,013,464 |
| Max. Reads per Primer | 2,013,464 |
| Min. Reads per Primer | 2,013,464 |
| Reads without Primers | 166,327 |
| Percent Bases in Reads with Primers | 0.9243 |
| Percent Reads with Primers | 0.9237 |
| Number of High-Quality Isoforms | 122,253 |
| Number of Low-Quality Isoforms | 82 |

**Table S5.** Reference guided transcriptome assembly statistics for short read transcriptome data assembled with Stringtie and Scallop for FLNC reads assembled with StringTie.

| **Stat** | **Tim_Flg (StringTie)** | **Tim_Grn (StringTie)** | **Tim_RT (StringTie)** | **Tim_Spk (StringTie)** | **Tim_Ypl_am (StringTie)** | **Tim_Ypl_pm (StringTie)** | **Tim_Flg (Scallop)** | **Tim_Grn (Scallop)** | **Tim_RT (Scallop)** | **Tim_Spk (Scallop)** | **Tim_Ypl_am (Scallop)** | **Tim_Ypl_pm (Scallop)** | **FLNC (StringTie)** |
| --- | --- | --- | --- | --- | --- | --- | --- | --- | --- | --- | --- | --- | --- |
| **Number of genes** | 64,440 | 74,303 | 74,018 | 80,533 | 78,710 | 74,985 | 88,176 | 96,768 | 110,719 | 120,211 | 119,355 | 108,452 | 38,266 |
| **Number of Transcripts** | 107,001 | 116,848 | 122,246 | 131,678 | 121,371 | 119,890 | 171,587 | 178,280 | 207,036 | 219,391 | 199,944 | 194,294 | 47,645 |
| **Transcripts per gene** | 1.66 | 1.57 | 1.65 | 1.64 | 1.54 | 1.6 | 1.95 | 1.84 | 1.87 | 1.83 | 1.68 | 1.79 | 1.25 |
| **Number of monoexonic genes** | 9,042 | 8,029 | 11,696 | 13,082 | 16,772 | 12,699 | 9,722 | 9,324 | 16,010 | 18,183 | 28,058 | 16,834 | 2,267 |
| **Monoexonic transcripts** | 9,608 | 8,429 | 12,399 | 13,995 | 17,672 | 13,479 | 13,582 | 12,364 | 22,207 | 24,503 | 35,648 | 22,086 | 2,613 |
| **Transcript mean size cDNA (bp)** | 1,870.56 | 1,695.45 | 1,804.84 | 1,692.87 | 1,565.93 | 1,687.33 | 1,701.11 | 1,611.81 | 1,642.09 | 1,500.57 | 1,343.54 | 1,481.63 | 2,288.66 |
| **Transcript median size cDNA (bp)** | 1,589 | 1,458 | 1,555 | 1,458 | 1,370 | 1,458 | 1,412 | 1,348 | 1,364 | 1,243 | 1,104 | 1,230 | 2,083 |
| **Min cDNA** | 200 | 200 | 200 | 200 | 200 | 200 | 200 | 205 | 200 | 200 | 200 | 200 | 214 |
| **Max cDNA** | 29,570 | 24,804 | 22,841 | 22,800 | 18,140 | 24,471 | 27,601 | 24,173 | 19,679 | 22,899 | 19,721 | 24,230 | 11,557 |
| **Total exons** | 590,973 | 633,237 | 673,868 | 691,273 | 612,859 | 629,905 | 859,672 | 896,823 | 1,000,394 | 1,023,926 | 858,851 | 903,298 | 314,594 |
| **Exons per transcript** | 5.52 | 5.42 | 5.51 | 5.25 | 5.05 | 5.25 | 5.01 | 5.03 | 4.83 | 4.67 | 4.3 | 4.65 | 6.6 |
| **Exon mean size (bp)** | 338.68 | 312.85 | 327.42 | 322.47 | 310.12 | 321.15 | 339.53 | 320.41 | 339.84 | 321.52 | 312.78 | 318.69 | 346.62 |
| **Intron mean size (bp)** | 627.99 | 636.98 | 637.24 | 628.67 | 631.46 | 642.5 | 632.04 | 655.41 | 625.35 | 623.85 | 641.96 | 634.87 | 499.34 |

| **Sample_Code** | **Sample_Description** |
| --- | --- |
| Tim_Flg | Timopheevii Flag Leaf |
| Tim_Grn | Timopheevii Grains |
| Tim_RT | Timopheevii Roots am |
| Tim_Ypl_am | Timopheevii seedlings dawn |
| Tim_Ypl_pm | Timopheevii seedlings dusk |
| Tim_Spk | Timopheevii Spike |

**Table S6.** REAT Transcriptome Mikado consolidated gene sets, gene model statistics.

| **Stat** | **mikado_long.loci** | **mikado_all.loci** |
| --- | --- | --- |
| **Number of genes** | 38,734 | 76,872 |
| **Number of Transcripts** | 43,617 | 118,829 |
| **Transcripts per gene** | 1.13 | 1.55 |
| **Number of monoexonic genes** | 4,965 | 11,082 |
| **Monoexonic transcripts** | 5,192 | 12,238 |
| **Transcript mean size cDNA (bp)** | 2,105.83 | 1,894.44 |
| **Transcript median size cDNA (bp)** | 1,889 | 1,671 |
| **Min cDNA** | 402 | 200 |
| **Max cDNA** | 11,557 | 16,724 |
| **Total exons** | 271,900 | 655,747 |
| **Exons per transcript** | 6.23 | 5.52 |
| **Exon mean size (bp)** | 337.81 | 343.29 |
| **CDS mean size (bp)** | 257.15 | 242.85 |
| **Transcript mean size CDS (bp)** | 1,435.87 | 1,135.21 |
| **Transcript median size CDS (bp)** | 1,275 | 948 |
| **Min CDS** | 0 | 0 |
| **Max CDS** | 10,899 | 16,092 |
| **Intron mean size (bp)** | 469.06 | 505.98 |
| **5'UTR mean size (bp)** | 230.09 | 270.55 |
| **3'UTR mean size (bp)** | 373.81 | 419.66 |

**Table S7.** List of Species used for cross species protein alignment.

| **NCBI_RefSeq_ID** | **Species** |
| --- | --- |
| GCF_000003195.3 | *Sorghum bicolor* |
| GCF_000005505.3 | *Brachypodium distachyon* |
| GCF_000263155.2 | *Setaria italica* |
| GCF_001433935.1 | *Oryza sativa* |
| GCF_002162155.2 | *Triticum dicoccoides* |
| GCF_002211085.1 | *Panicum hallii* |
| GCF_002575655.2 | *Aegilops tauschii subsp. strangulata* |
| GCF_016808335.1 | *Panicum virgatum* |
| GCF_902167145.1 | *Zea mays* |
| GCF_904849725.1 | *Hordeum vulgare* |

**Table S8.** BUSCO evaluation results of (a) genome assembly and (b) gene model prediction

|  | **(a) Genome assembly** | | **(b) Gene model prediction** | |
| --- | --- | --- | --- | --- |
|  | Value | % | Value | % |
| **Complete BUSCOs** | 1596 | 98.9 | 1609 | 99.7 |
| **Complete and single-copy BUSCOs** | 172 | 10.7 | 32 | 2.0 |
| **Complete and duplicated BUSCOs** | 1424 | 88.2 | 1577 | 97.7 |
| **Fragmented BUSCOs** | 9 | 0.6 | 1 | 0.1 |
| **Missing BUSCOs** | 9 | 0.5 | 4 | 0.2 |
| **Total BUSCO groups searched** | 1614 | | 1614 | |
